# Genomic characterization and computational phenotyping of nitrogen-fixing bacteria isolated from Colombian sugarcane fields

**DOI:** 10.1101/780809

**Authors:** Luz K. Medina-Cordoba, Aroon T. Chande, Lavanya Rishishwar, Leonard W. Mayer, Lina C. Valderrama-Aguirre, Augusto Valderrama-Aguirre, John Christian Gaby, Joel E. Kostka, I. King Jordan

**Affiliations:** School of Biological Sciences, Georgia Institute of Technology, Atlanta, Georgia, USA; PanAmerican Bioinformatics Institute, Cali, Valle del Cauca, Colombia; Applied Bioinformatics Laboratory, Atlanta, Georgia, USA; Laboratory of Microorganismal Production (Bioinoculums), Department of Field Research in Sugarcane, INCAUCA S.A.S., Cali, Valle del Cauca, Colombia; Biomedical Research Institute (COL0082529), Cali, Valle del Cauca, Colombia; Universidad Santiago de Cali, Cali, Colombia; Faculty of Chemistry, Biotechnology and Food Science, Norwegian University of Life Sciences, Ås, Norway

**Keywords:** biofertilizer, nitrogen fixation, plant growth promoter, genome sequencing, computational phenotyping

## Abstract

Previous studies have shown that the sugarcane microbiome harbors diverse plant growth promoting (PGP) microorganisms, including nitrogen-fixing bacteria, and the objective of this study was to design a genome-enabled approach to prioritize sugarcane associated nitrogen-fixing bacteria according to their potential as biofertilizers. Using a systematic high throughput approach, 22 pure cultures of nitrogen-fixing bacteria were isolated and tested for diazotrophic potential by PCR amplification of nitrogenase (*nifH*) genes, common molecular markers for nitrogen fixation capacity. Genome sequencing confirmed the presence of intact nitrogenase *nifH* genes and operons in the genomes of 18 of the isolates. Isolate genomes also encoded operons for phosphate solubilization, siderophore production operons, and other PGP phenotypes. *Klebsiella pneumoniae* strains comprised 14 of the 22 nitrogen-fixing isolates, and four others were members of closely related genera to *Klebsiella*. A computational phenotyping approach was developed to rapidly screen for strains that have high potential for nitrogen fixation and other PGP phenotypes while showing low risk for virulence and antibiotic resistance. The majority of sugarcane isolates were below a genotypic and phenotypic threshold, showing uniformly low predicted virulence and antibiotic resistance compared to clinical isolates. Six prioritized strains were experimentally evaluated for PGP phenotypes: nitrogen fixation, phosphate solubilization, and the production of siderophores, gibberellic acid and indole acetic acid. Results from the biochemical assays were consistent with the computational phenotype predictions for these isolates. Our results indicate that computational phenotyping is a promising tool for the assessment of benefits and risks associated with bacteria commonly detected in agricultural ecosystems.

**IMPORTANCE:** A genome-enabled approach was developed for the prioritization of native bacterial isolates with the potential to serve as biofertilizers for sugarcane fields in Colombia’s Cauca Valley. The approach is based on computational phenotyping, which entails predictions related to traits of interest based on bioinformatic analysis of whole genome sequences. Bioinformatic predictions of the presence of plant growth promoting traits were validated with experimental assays and more extensive genome comparisons, thereby demonstrating the utility of computational phenotyping for assessing the benefits and risks posed by bacterial isolates that can be used as biofertilizers. The quantitative approach to computational phenotyping developed here for the discovery of biofertilizers has the potential for use with a broad range of applications in environmental and industrial microbiology, food safety, water quality, and antibiotic resistance studies.

## INTRODUCTION

The human population is expected to double in size within the next 50 years, which will in turn lead to a massive increase in the global demand for food (1). Given the scarcity of arable land worldwide, an increase in agricultural production of this magnitude will require vast increases in cropping intensity and yield (2). It has been estimated that as much as 90% of the increase in global crop production will need to come from increased yield alone (3). At the same time, climate change and other environmental challenges will necessitate the development of agricultural practices that are more ecologically friendly and sustainable.

Chemical fertilizers that provide critical macronutrients to crops – such as nitrogen (N), phosphorus (P), potassium (K), and sulfur (S) – are widely used to maximize agricultural yield (4). The application of chemical fertilizers represents a major cost for agricultural companies and also contributes to environmental damage, in the form of eutrophication, hypoxia, harmful algal blooms, and air pollution through the formation of microparticles (5). Biological fertilizers (biofertilizers) are comprised of microbial inoculants that promote plant growth, thereby representing an alternative or complementary approach for increasing crop yield, which is more sustainable and environmentally friendly. Biofertilizers augment plant growth through nutrient acquisition, hormone production, and by boosting immunity to pathogens (6).

Sugarcane is a tall, perennial grass cultivated in tropical and warm temperate regions around the world, which is capable of producing high concentrations of sugar (sucrose) and diverse byproducts (7). Sugarcane is consistently ranked as one of the top ten planted crops in the world (8). Sugarcane agriculture plays a vital role in the economy of Colombia by supporting the production of food products and biofuel (ethanol). The long-term goals of this work are to develop more effective and sustainable sugarcane cropping practices in Colombia by simultaneously (i) increasing crop yield, and (ii) decreasing the reliance on chemical fertilizers via the discovery, characterization, and application of endemic (native) biofertilizers to Colombian sugarcane fields.

Most sugarcane companies in Colombia currently use commercially available biofertilizers, consisting primarily of nitrogen-fixing bacteria, which were discovered and isolated from other countries (primarily Brazil), with limited success. We hypothesized that indigenous bacteria should be better adapted to the local environment and thereby serve as more effective biofertilizers for Colombian sugarcane. The use of indigenous bacteria as biofertilizers should also mitigate potential threats to the environment posed by non-native, and potentially invasive, species of bacteria. Finally, indigenous bacteria represent a renewable resource that agronomists can continually develop through isolation and cultivation of local strains.

The advent of next-generation sequencing technologies has catalyzed the development of genome-enabled approaches to harness plant microbiomes in sustainable agriculture (9, 10). The objective of this study was to use genome analysis to predict the local bacterial isolates that have the greatest potential for plant growth promotion while representing the lowest risk for virulence and antibiotic resistance. Putative biofertilizer strains were isolated and cultivated from Colombian sugarcane fields, and computational phenotyping was employed to predict their potential utility as biofertilizers. We then performed a laboratory evaluation of the predicted plant growth promoting properties of the prioritized bacterial biofertilizer isolates, with the aim of validating our computational phenotyping approach.

## RESULTS

### Initial genome characterization of putative nitrogen-fixing bacteria

A systematic cultivation approach, incorporating seven carbon substrates in nitrogen-free media (Fig. S1), was employed to isolate putative nitrogen-fixing bacteria from four different sugarcane plant compartments, and isolates were screened for nitrogen fixation potential through PCR amplification of *nifH* genes. This initial screening procedure yielded several hundred clonal isolates of putative nitrogen-fixing bacteria, and Ribosomal Intergenic Spacer Analysis (RISA) was subsequently used to identify the (presumably) genetically unique strains from the larger set of clonal isolates. A total of 22 potentially unique strains of putative nitrogen-fixing bacteria were isolated in this way and selected for genome sequence analysis.

Genome sequencing and assembly summary statistics for the 22 isolates are shown in Table 1. Isolate genomes were sequenced to an average of 67x coverage (range: 50x – 88x) and genome sizes range from 4.5Mb to 6.1Mb. GC content varies from 41.82% – 66.69%, with a distinct mode at ∼57%. The genome assemblies are robust with a range of 24 – 294 contigs ≥500bp in length and averages of N50=310,166bp and L50=8.4. Genome sequence assemblies, along with their functional annotations, can all be found using the NCBI BioProject PRJNA418312. Individual BioSample, Genbank Accession, and Assembly Accession numbers for the 22 isolates are shown in Table S1.

**Table 1.** Genome assembly statistics for the isolates characterized here.

| Sample ID | Genome Length<br>(bp) | N50<sup>a</sup> | L50<sup>b</sup> | GC(%) | # of<br>Contigs<sup>c</sup> |
| --- | --- | --- | --- | --- | --- |
| SCK1 | 4,522,541 | 402,304 | 4 | 66.79 | 24 |
| SCK2 | 5,231,439 | 417,927 | 5 | 59.33 | 53 |
| SCK3 | 3,824,428 | 670,745 | 3 | 41.82 | 150 |
| SCK4 | 4,511,030 | 223,239 | 8 | 66.79 | 55 |
| SCK5 | 5,774,634 | 162,673 | 13 | 53.1 | 98 |
| SCK6 | 6,094,823 | 117,689 | 15 | 56.73 | 294 |
| SCK7 | 5,693,007 | 282,996 | 7 | 57.03 | 50 |
| SCK8 | 5,695,902 | 281,292 | 9 | 57.03 | 50 |
| SCK9 | 5,579,618 | 311,650 | 6 | 57.03 | 42 |
| SCK10 | 5,591,472 | 614,324 | 3 | 57.03 | 34 |
| SCK11 | 5,696,136 | 382,597 | 5 | 57.15 | 268 |
| SCK12 | 5,817,089 | 176,655 | 10 | 57.02 | 79 |
| SCK13 | 5,476,221 | 358,490 | 5 | 57.34 | 33 |
| SCK14 | 5,465,811 | 300,899 | 5 | 57.34 | 41 |
| SCK15 | 5,564,330 | 330,579 | 5 | 57.15 | 43 |
| SCK16 | 5,795,921 | 478,592 | 3 | 54.06 | 84 |
| SCK17 | 5,475,984 | 358,490 | 4 | 57.34 | 35 |
| SCK18 | 5,476,135 | 422,400 | 3 | 57.34 | 32 |
| SCK19 | 5,688,396 | 270,585 | 7 | 57.09 | 56 |
| SCK20 | 5,500,801 | 82,111 | 20 | 57.45 | 165 |
| SCK21 | 5,324,920 | 112,078 | 15 | 55.26 | 100 |
| SCK22 | 5,847,607 | 65,329 | 29 | 57.02 | 181 |

<sup>a</sup> When the contigs of an assembly are arranged from largest to smallest, N50 is the length of the contig that makes up at least 50% of the genome
<sup>b</sup> L50 is the number of contigs equal to or longer than N50 In other words, L50, for example, is the minimal number of contigs that cover half the assembly
<sup>c</sup> Number of contigs $\geq 500$ bp in length

### Comparative genomic analysis

Average nucleotide identity (ANI; Fig. 1) and 16S rRNA gene sequence analysis (Fig. S2) were employed in the taxonomic assignment of nitrogen-fixing isolates and the results of both approaches were highly concordant (Table 2), with ANI yielding superior resolution to 16S rRNA gene sequence analysis. A total of eight different species and seven different genera were identified among the 22 isolates characterized. Analysis of *nifH* gene sequences also gave similar results; however, four of the isolates were not found to encode *nifH* genes, despite their (apparent) ability to grow on nitrogen-free media and the positive *nifH* PCR results. This could be due to false-positives in the original PCR analysis for the presence of *nifH* genes, or to changes in the composition of (possibly mixed) bacterial cultures during subsequent growth steps after the initial isolation on nitrogen-free media.

**FIG 1.**
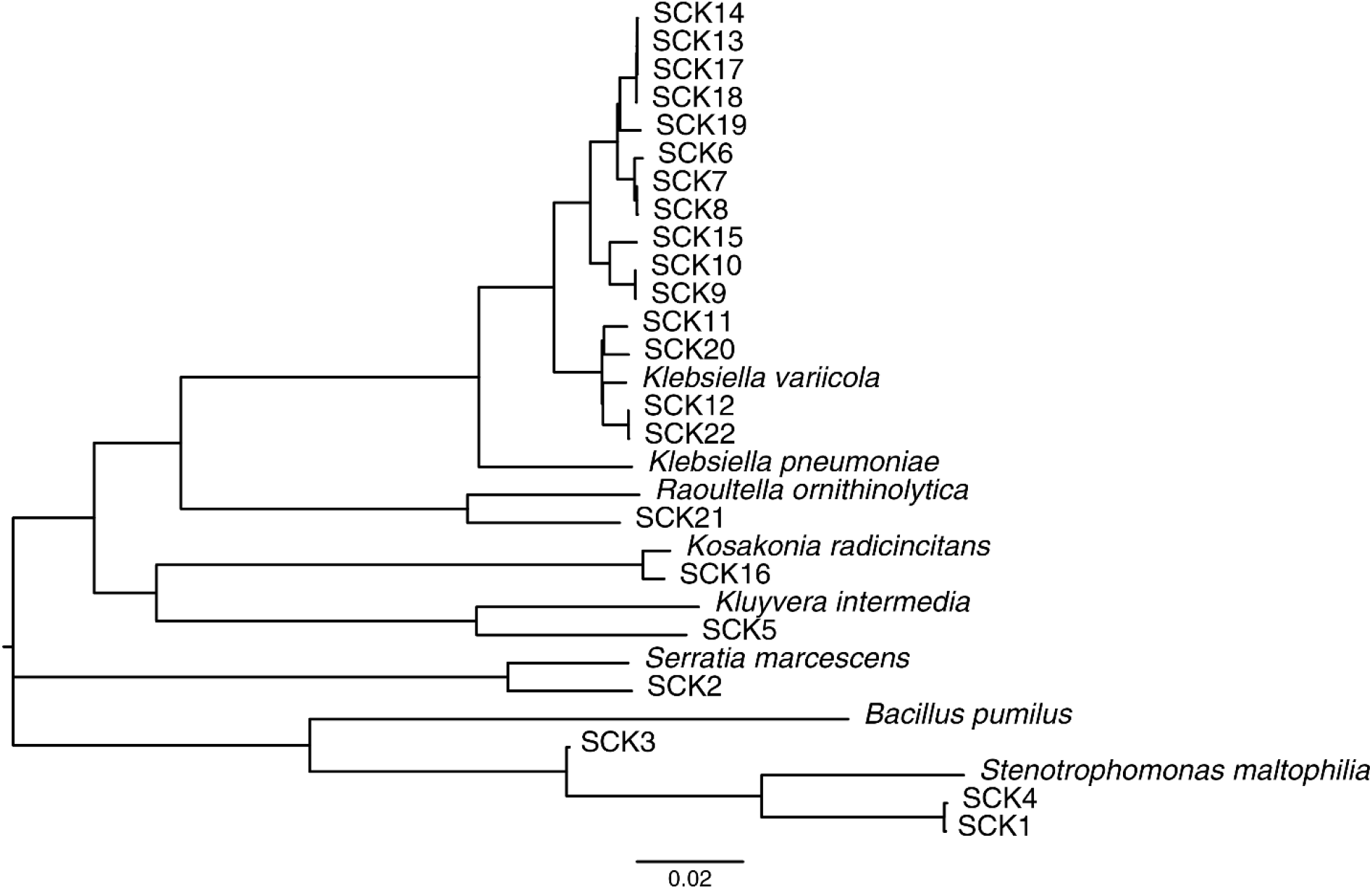
Phylogeny of the bacterial isolates characterized here (SCK numbers) together with their most closely related bacterial type strains. The phylogeny was reconstructed using pairwise average nucleotide identities between whole genome sequence assemblies, converted to p-distances, with the neighbor-joining method. Horizontal branch lengths are scaled according the p-distances as shown.

**Table 2.** Identity of the most closely related species (genus) for the isolates characterized here. Species (genus) identification was performed using average nucleotide identity (ANI), 16S rRNA and *nifH* sequence comparisons.

| Strain | ANI | 16S | *nifH* |
| --- | --- | --- | --- |
| SCK1 | *Stenotrophomonas maltophilia* | *Stenotrophomonas* | NA |
| SCK2 | *Serratia marcescens* | *Serratia* | NA |
| SCK3 | *Bacillus pumilus* | *Bacillus* | NA |
| SCK4 | *Stenotrophomonas maltophilia* | *Stenotrophomonas* | NA |
| SCK5 | *Kluyvera intermedia* | *Kluyvera* | *Kluyvera* |
| SCK6 | *Klebsiella pneumoniae* | *Klebsiella* | *Klebsiella* |
| SCK7 | *Klebsiella pneumoniae* | *Klebsiella* | *Klebsiella* |
| SCK8 | *Klebsiella pneumoniae* | *Klebsiella* | *Klebsiella* |
| SCK9 | *Klebsiella pneumoniae* | *Klebsiella* | *Klebsiella* |
| SCK10 | *Klebsiella pneumoniae* | *Klebsiella* | *Klebsiella* |
| SCK11 | *Klebsiella pneumoniae* | *Klebsiella* | *Klebsiella* |
| SCK12 | *Klebsiella pneumoniae* | *Klebsiella* | *Klebsiella* |
| SCK13 | *Klebsiella pneumoniae* | *Klebsiella* | *Klebsiella* |
| SCK14 | *Klebsiella pneumoniae* | *Klebsiella* | *Klebsiella* |
| SCK15 | *Klebsiella pneumoniae* | *Klebsiella* | *Klebsiella* |
| SCK16 | *Kosakonia radicincitans* | *Kosakonia* | *Kosakonia* |
| SCK17 | *Klebsiella pneumoniae* | *Klebsiella* | *Klebsiella* |
| SCK18 | *Klebsiella pneumoniae* | *Klebsiella* | *Klebsiella* |
| SCK19 | *Klebsiella pneumoniae* | *Klebsiella* | *Klebsiella* |
| SCK20 | *Klebsiella pneumoniae* | *Klebsiella* | *Klebsiella* |
| SCK21 | *Raoultella ornithinolytica* | *Raoultella* | *Raoultella* |
| SCK22 | *Klebsiella variicola* | *Klebsiella* | *Klebsiella* |

The majority of isolates, 14 of 22, were characterized as *Klebsiella pneumoniae*, consistent with previous studies showing that *K. pneumoniae* strains are capable of fixing nitrogen (11); in fact, the canonical *nif* operons were defined in the *K. pneumoniae* type strain 342 genome sequence (12). *K. pneumoniae* is also known to be an opportunistic pathogen that can cause disease in immunocompromised human hosts (13), which raises obvious safety concerns regarding its application to crops as part of a biofertilizer inoculum. We performed a comparative sequence analysis between the endophytic nitrogen-fixing *K. pneumoniae* type strain 342, which is capable of infecting the mouse urinary tract and lung (14), and five of the isolates identified as *K. pneumoniae* here. All genomes were shown to contain the *nif* cluster, which contains five functionally related *nif* operons involved in nitrogen fixation (Fig. 2). In contrast, the four most critical pathogenicity islands implicated in the virulence of *K. pneumoniae* 342 were all missing in the environmental *K. pneumoniae* isolates characterized here (PAI 1-4 in Fig. 2A). The absence of pathogenicity islands in the genome of the endophytic nitrogen-fixer *K. michiganensis* Kd70 was associated with an inability to infect the urinary tract in mice (15). Our results indicate that nitrogen-fixing *K. pneumoniae* environmental isolates from Colombian sugarcane fields do not pose a health risk compared to clinical and environmental isolates that have previously been associated with pathogenicity. We explore this possibility in more detail in the following section on computational phenotyping.

**FIG 2.**
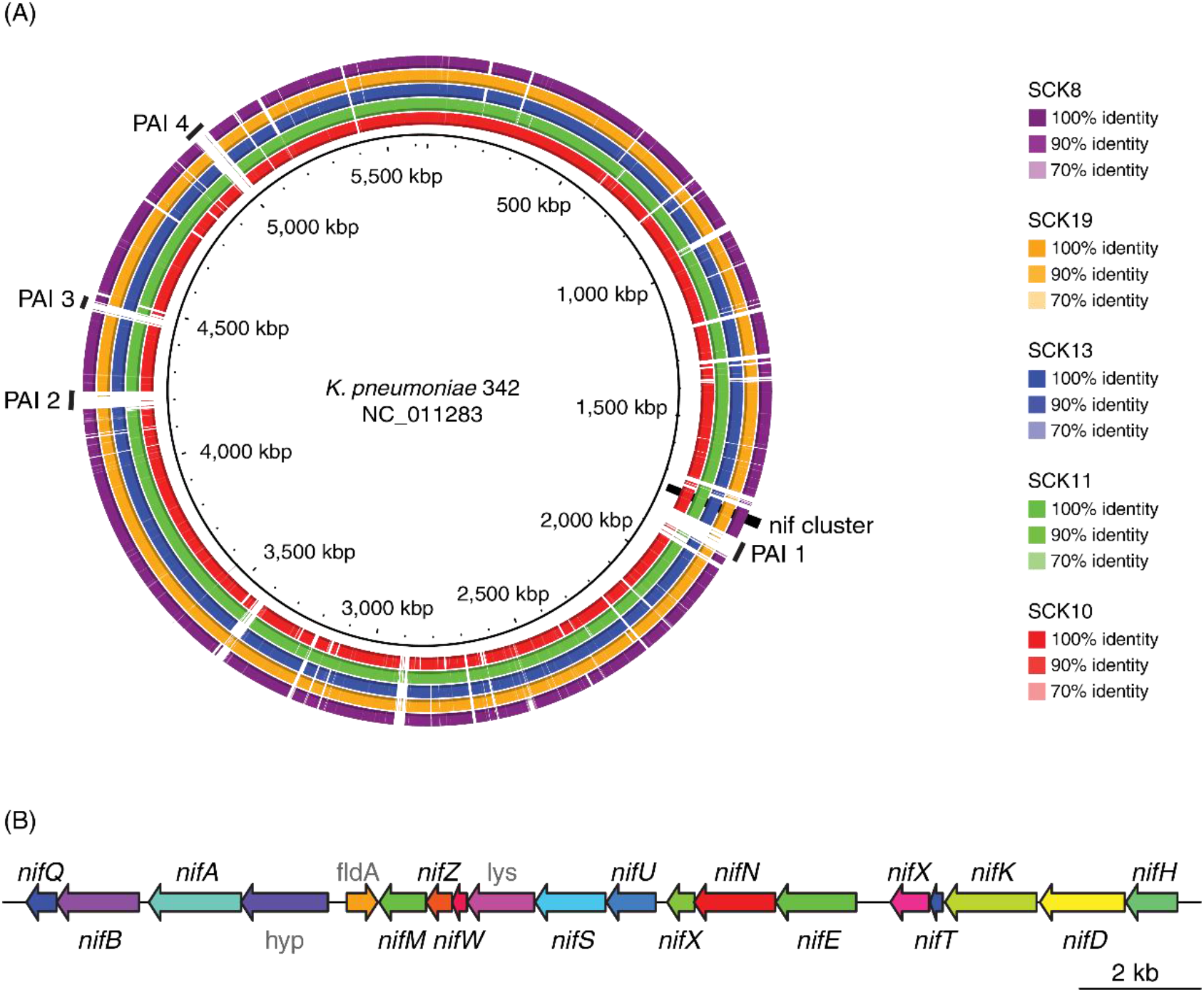
Comparison of the *K. pneumoniae* type strain 342 to *K. pneumoniae* sugarcane isolates characterized here. (A) BLAST ring plot showing synteny and sequence similarity between *K. pneumoniae* 342 and five *K. pneumoniae* sugarcane isolates. The *K. pneumoniae* 342 genome sequence is shown as the inner ring, and syntenic regions of the five *K. pneumoniae* sugarcane isolates are shown as rings with strain-specific color-coding according to the percent identity between regions of *K. pneumoniae* 342 and the sugarcane isolates. The genomic locations of *nif* operon cluster along with four important pathogenicity islands (PAIs) are indicated. PAI1 – type IV secretion and aminoglycoside resistance, PAI2 hemolysin and fimbria secretion, heme scavenging, PAI3 – radical S-adenosyl-L-methionine (SAM) and antibiotic resistance pathways, PAI4 – fosfomycin resistance and hemolysin production. (B) A scheme of the *nif* operon cluster present in both *K. pneumoniae* 342 and the five *K. pneumoniae* sugarcane isolates.

The *nifH* genes from the *Klebsiella* isolates characterized here form two distinct phylogenetic clusters (Fig. 3). This finding is consistent with previous results showing multiple clades of *nifH* among *Klebsiella* genome sequences (16–18) and underscores the potential functional diversity, with respect to nitrogen fixation, for the sugarcane isolates.

**FIG 3.**
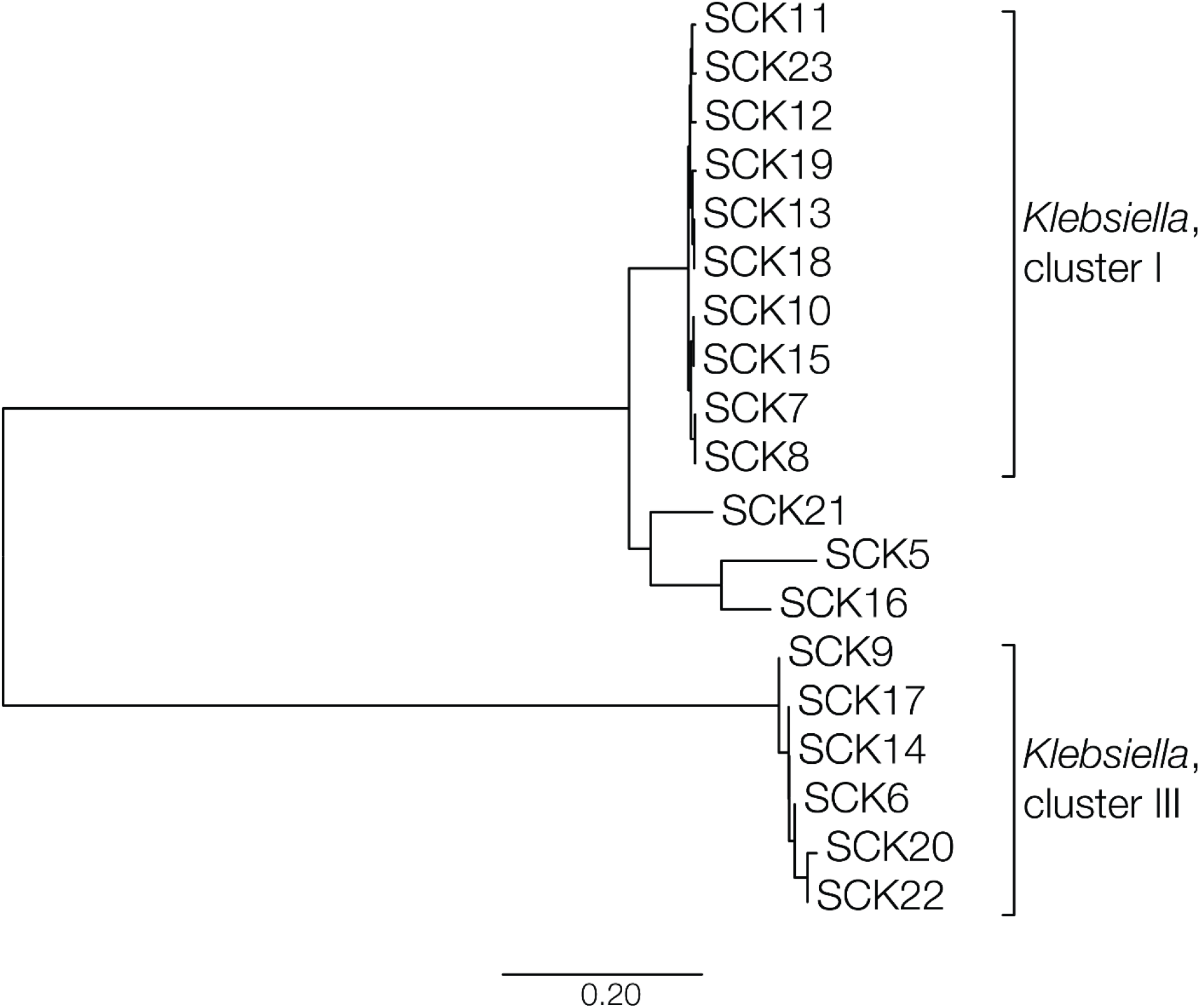
Phylogeny of the *nifH* genes for the *Klebsiella* bacterial isolates characterized here (SCK numbers). The phylogeny was reconstructed using pairwise nucleotide p-distances between *nifH* genes recovered from the isolate genome sequences using the neighbor-joining method. Horizontal branch lengths are scaled according the p-distances as shown.

### Computational phenotyping

Computational phenotyping, also referred to as reverse genomics, was used to evaluate the potential of the bacterial isolates characterized here to serve as biofertilizers for Colombian sugarcane fields. For the purpose of this study, computational phenotyping entails the prediction of specific organismal phenotypes, or biochemical capacities, based on the analysis of functionally annotated genome sequences (19). The goal of the computational phenotyping performed here was to identify isolates that show the highest predicted capacity for plant growth promotion while presenting the lowest risk to human populations. Accordingly, bacterial isolate genome sequences were screened for gene features that correspond to the desirable (positive) characteristics of (i) nitrogen fixation and (ii) plant growth promotion and the disadvantageous (negative) characteristics of (iii) virulence and (iv) antimicrobial resistance. Genome sequences were scored and ranked according to the combined presence or absence of these four categories of gene features as described in the Materials and Methods. To compute genome scores, the presence of nitrogenase and plant growth promoting genes contribute positive values, whereas the presence of virulence factors and predicted antibiotic resistance yield negative values. Scores for each of the four specific phenotypic categories were normalized and combined to yield a single composite score for each bacterial isolate genome. The highest scoring isolates are predicted as best candidates to be included as part of a sugarcane biofertilizer inoculum (Figure 4; Table S2). The predicted biochemical capacities of the highest scoring isolates were subsequently experimentally validated.

**FIG 4.**
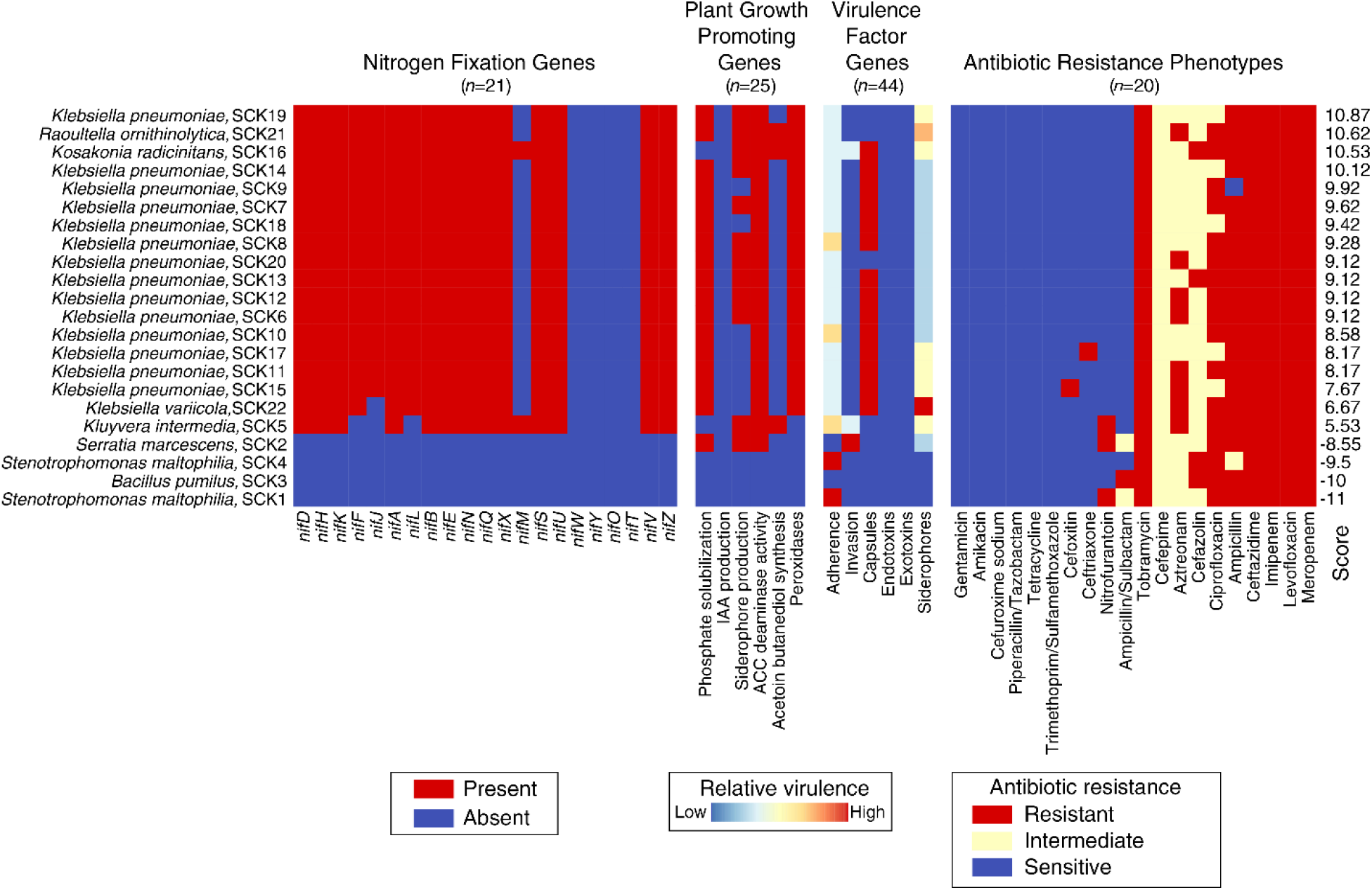
Computational phenotyping of the sugarcane bacterial isolates characterized here. The presence (red) and absence (blue) profiles for nitrogen fixation genes, plant growth promoting genes, and virulence factor genes are shown for the 22 bacterial isolates. Results are shown for all *n*=21 nitrogen-fixing genes. Results for plant growth promoting genes (*n*=25) and virulence factor genes (*n*=44) are merged into six gene categories each. Predicted antibiotic resistance profiles are shown for *n*=20 antibiotic classes. Detailed results for gene presence/absence and predicted antibiotic resistance profiles are shown in Table S2. The results for all four phenotypic classes of interest were merged into a single priority score for each isolates (right side of plot), as described in the Materials and Methods, and used to rank the isolates with respect to their potential as biofertilizers.

Isolates are ranked according to their composite genome scores, with a value of 10.87 observed as the highest potential for biofertilizer production (Figure 4). Individual gene and phenotype scores are color coded for each genome, and the four functional-specific categories are shown separately. The *nif* gene presence/absence profiles were found to be highly similar for all but four of the bacterial isolates characterized here, those which are not members of the *Klebsiella* genus, or closely related species, and do not encode any *nif* genes. The four non-nitrogen fixing isolates represent bacterial species that are commonly found in soil (20–23), but they are not predicted to be viable biofertilizers. The *Kosakonia radicincitans* genome encodes the largest number of *nif* genes (*n*=17) observed for any of the Colombian sugarcane isolates. This is consistent with previous studies showing that isolates of this species are capable of fixing nitrogen (24). The 14 characterized *K. pneumoniae* genomes all contain 16 out of 21 *nif* genes, including the core *nifD* and *nifK* genes, which encode the heterotetramer core of the nitrogenase enzyme, and the *nifH* gene, which encodes the dinitrogenase reductase subunit (25). These genomes also all encode the nitrogenase master regulators *nifA* and *nifL*. The missing *nif* genes for the *K. pneumoniae* isolates correspond to accessory structural and regulatory proteins that are not critical for nitrogen fixation. Accordingly, all of *K. pneumoniae* isolate genomes are predicted to encode the capacity for nitrogen fixation, consistent with previous results (14, 26). The single *Raoultella ornithinolytica* isolate characterized here also contains the same 16 *nif* genes; *Raoultella* species have previously been isolated from sugarcane (27) and have also been demonstrated to fix nitrogen (28).

Initially, a total of 29 canonical bacterial plant growth promoting genes were mined from the literature, 25 of which were found to be present in at least one of the bacterial isolate genome sequences characterized here. These 25 plant growth promoting genes were organized into six distinct functional categories: phosphate solubilization, indolic acetic acid (IAA) production, siderophore production, 1-aminocyclopropane-1-carboxylate (ACC) deaminase, acetoin butanediol synthesis, and peroxidases (Table S3). For the purposes of visualization (Fig. 4), each functional category is deemed to be present in an isolate genome sequence if all required genes for that function can be found, but the weighted scoring for these categories is based on individual gene counts as described in the Materials and Methods. The *R. ornithinolytica* isolate shows the highest predicted capacity for plant growth promotion, with 5 of the 6 functional categories found to be fully present. The majority of *K. pneumoniae* isolates also show similar, but not identical, plant growth promoting gene presence/absence profiles, with 3 or 4 functional categories present. The capacity for siderophore production is predicted to vary among *K. pneumoniae* isolates. The *K. radicincitans* genome also encodes 4 functional categories of plant growth promoting genes, but differs from the *K. pneumoniae* isolates with respect to absence of phosphate solubilization genes and the presence of acetoin butanediol synthesis genes. Three of the four species found to lack *nif* genes also do not score present for any of the plant growth promoting gene categories, further underscoring their predicted lack of utility as biofertilizers.

Initially, a total of ∼2,500 virulence factor genes were mined from the Virulence Factor Database (VFDB) (29), 44 of which were found to be present in at least one of the bacterial isolate genome sequences characterized here. These 44 virulence factors were organized into six distinct functional categories related to virulence and toxicity: adherence, invasion, capsules, endotoxins, exotoxins, and siderophores. The weighted scores for these categories were computed based on individual gene presence/absence patterns (Fig. 4). In contrast to the *K. pneumoniae* clinical isolates which have previously been characterized as opportunistic pathogens, the *K. pneumoniae* environmental isolates showed uniformly low virulence scores. The virulence factor genes found among the *K. pneumoniae* environmental isolates correspond to adherence proteins, capsules, and siderophores. As shown in Fig. 2, genomes of environmental isolates lack coding capacity for important invasion and toxin proteins, including the Type IV secretion system, which are found in clinical *K. pneumoniae* isolates. The *R. ornithinolytica* and *K. radicincitans* isolates, both of which show high scores for nitrogen fixation and plant growth promotion, gave higher virulence scores in comparison to the environmental *K. pneumoniae* isolates. Whereas *Bacillus pumilus* had the lowest virulence score for any of the isolates, the remaining three non-nitrogen fixing isolates had the highest virulence scores and were shown to encode well-known virulence factors, such as Type IV, hemolysin, and fimbria secretion systems.

The predicted antibiotic resistance phenotypes for all characterized isolates were fairly similar across the 20 classes of antimicrobial compounds for which predictions were made. The majority of the *K. pneumoniae* genomes, along with the relatively high scoring *R. ornithinolytica* and *K. radicincitans* isolate genomes, indicated predicted susceptibility to 10 of the 20 classes of antimicrobial compounds, intermediate susceptibility for 2-4, and predicted resistance to 5-8. The highest level of predicted antibiotic resistance was seen for *Serratia marcescens*, with resistance predicted for 8 compounds and intermediate susceptibility predicted for 4.

Computational phenotyping scores for the four categories were normalized and combined into a final score, with respect to their potential as biofertilizers (Fig. 4). Most of the top positions are occupied by *K. pneumoniae* isolates, with the exception of the second-ranked *R. ornithinolytica* and the third-ranked *K. radicincitans*. The results of a similar analysis of four additional plant associated *Klebsiella* genomes are shown in Fig. S3.

### Virulence comparison

The results described in the previous section indicate that the majority of the *K. pneumoniae* strains isolated from Colombian sugarcane fields have the highest overall potential as biofertilizers, including a low predicted potential for virulence. Nevertheless, the fact that strains of *K. pneumoniae* have previously been characterized as opportunistic pathogens (30) raises concerns when considering the use of *K. pneumoniae* as part of a bioinoculum that will be applied to sugarcane fields. With this in mind, we performed a broader comparison of the predicted virulence profiles for Colombian sugarcane isolates along with a collection of 28 clinical isolates of *K. pneumoniae* and several other closely related species (See Table S5 for isolate accession numbers). For this comparison, the same virulence factor scoring scheme described in the previous section was applied to all 50 genome sequences (Fig. 5). Perhaps most importantly, a very clear distinction was observed in the virulence score distribution, whereby all 28 clinical strains show a substantially higher predicted virulence (from 4.45 to 2.11) in comparison to the environmental isolates (1.55 to 0.00). Furthermore, the three environmental isolates that show the highest predicted virulence correspond to species with low predicted capacity for both nitrogen fixation and plant growth promotion; as such, these isolates would not be considered as potential biofertilizers. In particular, the *K. pneumoniae* environmental isolates showed uniformly low predicted virulence compared to clinical isolates of the same species. Thus, the results support, in principle, the use of the environmental *K. pneumoniae* isolates as biofertilizers for Colombian sugarcane fields.

**FIG 5.**
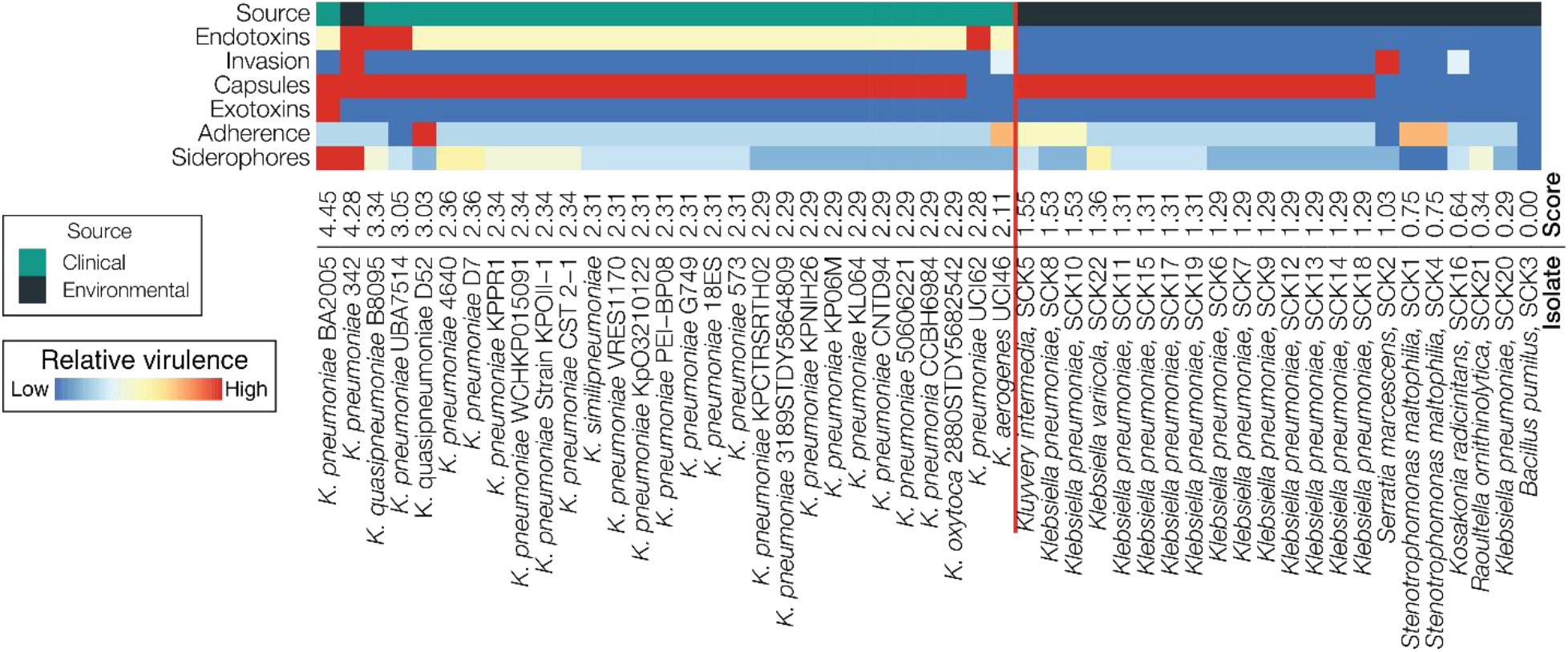
Comparison of predicted virulence profiles for clinical *K. pneumoniae* isolates compared to the environmental (sugarcane) bacterial isolates characterized here. As in Fig. 4, predicted virulence profiles for six classes of virulence factor genes are shown for each isolate. Isolate-specific virulence factor scores are shown for each isolate are based on the presence/absence profiles for the *n*=44 virulence factor genes as described in the Materials and Methods. The virulence factor genes are used to rank the genomes from most (left) to least (right) virulent. Clinical versus environmental samples are shown to the left and right, respectively, of the red line, based on their virulence scores.

### Experimental validation of prioritized isolates

The top six scoring isolates from the computational phenotyping were subjected to a series of cultivation-based phenotypic assays in order to validate their predicted biochemical activities: (i) acetylene reduction (a proxy for nitrogen fixation), (ii) phosphate solubilization, (iii) siderophore production, (iv) gibberellic acid production, and (v) indole acetic acid production.

Nitrogen fixation activity, as determined by acetylene reduction to ethylene, was observed in all six isolates, three of which had higher levels in comparison to the positive control (Fig. 6A). All six of the isolates showed high levels of phosphate solubilization (Fig. 6B & C) and siderophore production (Fig. 6D & E) compared to the respective negative controls. All six isolates showed the ability to produce gibberellic acid (Fig. 6F), whereas none were able to produce indole acetic acid. The biochemical assay results are consistent with the computational phenotype predictions for these isolates.

**FIG 6.**
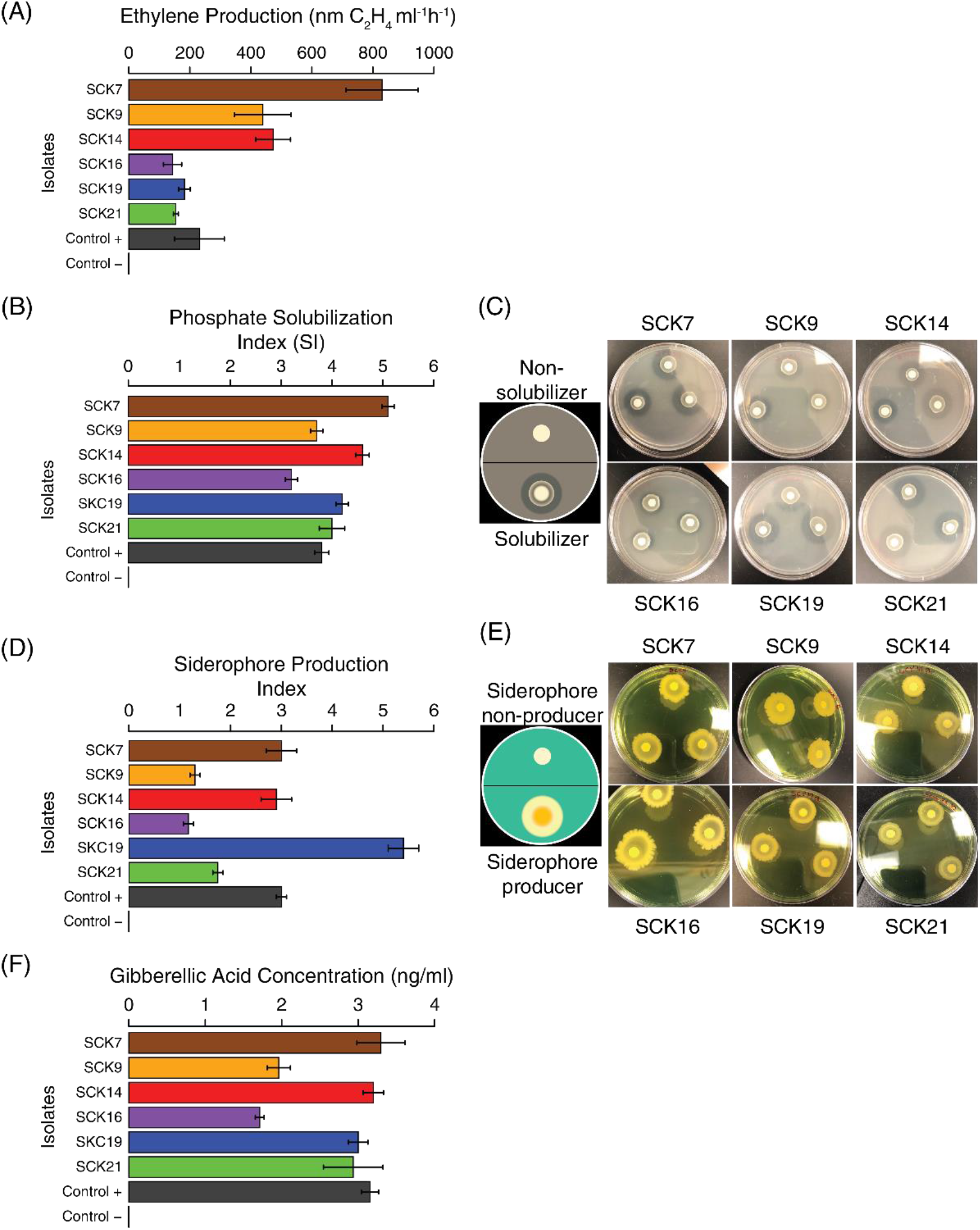
Experimental validation of prioritized biofertilizer isolates. The computationally predicted plant growth promoting phenotypes for the top six isolates were experimentally validated. All six strains were capable of acetylene reduction, i.e. ethylene production (A), phosphate solubilization (B&C), siderophore production (D&E), and gibberellic Acid production (F).

## DISCUSSION

Members of the *Enterobacteriaceae* are often observed in cultivation-independent studies of sugarcane and nitrogen-fixing *Enterobacteriaceae* are often isolated from sugarcane plants worldwide (31–35). The majority of isolates that were obtained in this study from Colombian sugarcane belonged to the family *Enterobacteriaceae*, with the *Klebsiella* as the most abundant genus along with *Serratia*, *Kluyvera, Stenotrophomonas*, and *Bacillus*. *Klebsiella* are Gram-negative, facultatively anaerobic bacteria found in soils, plants, or water (36). *Klebsiella* species have been isolated from a large variety of crops worldwide, such as sugarcane, rice, wheat, and maize (36–38). *Klebsiella* species associated with plants have been shown to fix nitrogen and express other plant growth promoting traits (37, 39). Specifically, *Klebsiella* species are abundant amongst the cultivable strains of *Enterobacteriaceae* obtained from sugarcane (31). For example, a survey of sugarcane in Guangxi, China observed that *Klebsiella* was the most abundant plant-associated nitrogen-fixing bacterial group (31), and among the strains isolated, *K. variicola* was shown to colonize sugarcane and promote plant growth (37). In addition, endophytic *Klebsiella* spp. have been isolated from commercial sugarcane in Brazil, and their potential for plant growth promotion was evaluated *in vitro* (40). Finally in Pakistan, the phenotypic diversity of plant growth promoting associated with sugarcane was determined, with *Klebsiella* also appearing as one of the most abundant bacteria found (33). At the same time, Klebsiella and other groups of *Enterobacteriaceae* commonly detected in agricultural systems are abundant in the human microbiome and often contain closely related members that are known opportunistic pathogens (41–44). The coexistence of microbial species that contain plant beneficial traits with closely related strains that potentially cause human diseases presents a challenge for the development of sustainable agriculture. How can we effectively perform a risk-benefit analysis of bacterial strains for potential use in the agricultural biotechnology industry? Thus, the overall goal of this study was to develop high throughput methods for the isolation and screening of nitrogen-fixing bacteria for their potential as biofertilizers.

### Computational phenotyping for the prioritization of potential biofertilizers

A computational phenotyping approach was developed for the screening of plant growth promoting bacteria for their potential to serve as biofertilizers. Computational phenotyping entails the implementation of a variety of bioinformatic and statistical methods to predict phenotypes of interest based on whole genome sequence analysis (45, 46). This approach has been used for a variety of applications in the biomedical sciences: prediction of clinically relevant phenotypes, study of infectious diseases, identification of opportunistic pathogenic bacteria in the human microbiome, and cancer treatment decisions (47, 48). To our knowledge, this study represents the first time computational phenotyping has been used for agricultural applications. To implement computational phenotyping for the prioritization of potential biofertilizers, we developed a scoring scheme based on the genome content of four functional gene categories of interest: nitrogen-fixing genes, other plant growth promoting genes, virulence factor genes, and antimicrobial resistance genes.

The results of the computational phenotyping predictions, confirmed by laboratory experiments, support the potential use of selected bacterial strains isolated from Colombian sugarcane fields as biofertilizers with minimum health risk to the human population. In particular, all isolates with higher scores (5.53 to 10.87, Fig. 4) in our scheme were found to demonstrate the potential to fix nitrogen and to promote plant growth in other ways, while lacking many of the important known virulence factors and antibiotic resistance genes that can be found in clinical isolates of the same species. In general, isolates SCK7, SCK14, and SCK19 appeared to possess more potent plant growth promoting properties compared to isolates SCK9, SCK16, and SCK21 (Fig. 4). Our computational phenotyping scheme also has valuable negative predictive value. Isolates that contained few or none of the beneficial traits that characterize biofertilizers, *Bacillus pumilus* SCK3 and *Stenotrophomonas maltophilia* SCK1, had the lowest scores (−10 and −11 respectively). Finally, it is also worth reiterating that the computationally predicted biochemical activities related to plant growth promotion were all validated by experimental results (Fig. 6).

### Virulence profiling for the prioritization of potential biofertilizers

Opportunistic pathogens are microorganisms that usually do not cause disease in a healthy host, but rather colonize and infect an immunocompromised host (49, 50). For example, *Klebsiella spp*. including *Klebsiella pneumoniae*, *Klebsiella oxytoca*, and *Klebsiella granulomatis* were associated with nosocomial diseases (51) and other hospital-acquired infections, primarily in immunocompromised persons (52). The potential for virulence, along with the presence of antimicrobial resistance genes, is an obvious concern when proposing to use *Klebsiella* spp. as biofertilizers. Importantly, we found that the environmental *Klebsiella* isolates did not contain pathogenicity islands associated with many virulence factor genes usually found in clinical isolates of *Klebsiella* spp. (Fig. 2). Our results are corroborated by a previous study of *Klebsiella michiganensis* Kd70 isolated from the intestine of larvae of *Diatraea saccharalis*, for which the genome was shown to contain multiple genes associated with plant growth promotion and root colonization, but lacked pathogenicity islands in its genome (15). In order to shed further light on this problem, we extended our study of environmental isolates from Colombian sugarcane to comparisons with genomes of *Klebsiella* clinical isolates associated with opportunistic infections in humans along with a number other environmental isolates with available genome sequences (Fig. 5). The virulence factor profiles for all of the environmental isolates were clearly distinct from the clinical strains, which show uniformly higher virulence profile scores, underscoring the relative safety of *Klebsiella* environmental isolates for use as biofertilizers.

### Potential for the use of computational phenotyping in other microbiology applications

The results obtained from the computational phenotyping approach developed in this study serve as a proof of principle in support of genomic guided approaches to sustainable agriculture. In particular, computational phenotyping can serve to substantially narrow the search space for potential plant growth promoting bacterial isolates, which can be further interrogated via experimental methods. Computational phenotyping can be used to simultaneously identify beneficial properties of plant associated bacterial isolates while avoiding potentially negative characteristics. In principle, this approach can be applied to a broad range of potential plant growth promoting isolates, or even assembled metagenomes, from managed agricultural ecosystems.

We can also envision a number of other potential applications for computational phenotyping of microbial genomes. The computational phenotyping methodology developed here has broad potential including diverse applications in agriculture, plant and animal breeding, food safety, water quality microbiology along with other industrial microbiology applications such as bioenergy, quality control/quality assurance, and fermentation microbiology as well as human health applications such as pathogen antibiotic resistance, virulence predictions, and microbiome characterization. For instance, computational phenotyping could be useful in food safety related to vegetable crop production. Vegetables harbor a diverse bacterial community dominated by the family *Enterobacteriaceae,* Gram-negative bacteria that include a huge diversity of plant growth promoting bacteria and enteric pathogens (53). Vegetables such as lettuce, spinach, and carrots are usually consumed raw, which increases the concern of bacterial infections or human disease outbreaks associated with consumption of vegetables (49).

Increasing antibiotic resistance, generated by the abuse of antibiotics in agriculture as well as medicine, is another major threat to human health (54), and the food supply chain creates a direct connection between the environmental habitat of bacteria and human consumers (55). Our computational phenotyping approach could provide for an additional food safety solution, which could be used to prevent the spread of antibiotic resistance pathogens genes present in the food chain.

## MATERIALS AND METHODS

### Sampling and cultivation of putative nitrogen-fixing bacteria from sugarcane

INCAUCA is a Colombian sugarcane company located in the Cauca River Valley in the southwest region of the country between the western and central Andes mountain ranges (http://www.incauca.com/). Samples of leaves, rhizosphere soil, stem, and roots were collected from the sugarcane fields 32T and 37T of the INCAUCA San Fernando farm located in the Cauca Valley (3°16’30.0“N 76°21’00.0”W). A high-throughput enrichment approach was developed to enable the cultivation of multiple strains of putative nitrogen-fixing bacteria from sugarcane field samples; details of this approach can be found in the Supplementary Material (Supplementary Methods and Fig. S1).

A total of 22 distinct *nifH* PCR+ isolates that passed the initial cultivation and screening steps were grown in LB medium (Difco) at 37°C for subsequent genomic DNA extraction. The E.Z.N.A. bacterial DNA kit (Omega Bio-Tek) was used for genomic DNA extraction, and paired-end fragment libraries (∼1,000bp) were constructed using the Nextera XT DNA library preparation kit (Illumina).

### Genome sequencing, assembly, and annotation

Isolate genomic DNA libraries were sequenced on the Illumina MiSeq platform using V3 chemistry, yielding approximately 400,000 paired-end 300bp sequence reads per sample. A list of all genome sequence analysis programs that were used for this study is provided Table S4. Sequence read quality control and trimming were performed using the programs FastQC version0.11.5 (56) and Trimmomatic (v.0.35) (57). *De novo* sequence assembly was performed using the program SPAdes (v.3.6) (58). Assembled genome sequences were annotated using the Rapid Annotations using Subsystems Technology (RAST) Web server (59, 60) and NCBI Prokaryotic Genome Annotation Pipeline (PGAP) (61). The 15 *Klebsiella* isolates characterized in this way were briefly described in a Genome Announcement (62), and the analysis here includes 7 additional non-*Klebsiella* isolates.

### Comparative genomic analysis

Average Nucleotide Identity (ANI) was employed to assign the taxonomy of the bacterial isolates characterized here (63, 64). Taxonomic assignment was also conducted by targeting small subunit ribosomal RNA (SSU rRNA) gene sequences. Nitrogenase enzyme encoding *nifH* gene sequences were extracted from isolate genome sequences, clustered, and taxonomically assigned using the TaxaDiva (v.0.11.3) method developed by our group (12). Whole genome sequence comparisons between bacterial isolates characterized here and the *K. pneumoniae* type strain 342 were performed using BLAST+ (v.2.2.28) (65) and visualized with the program CGView (v.1.0) (66). Details of the methods used comparative genomic analysis can be found in the Supplementary Methods section.

### Computational phenotyping

Computational phenotyping was performed by searching the bacterial isolate genome sequences characterized here for the presence/absence of genes or features related to four functional classes of interest, with respect to their potential as biofertilizers: (i) nitrogen fixation (NF), (ii) plant growth promotion (PGP), (iii) virulence factors (iv), and (4) antimicrobial resistance (AMR). Gene panels were manually curated by searching the literature (NCBI PubMed) for genes implicated in nitrogen fixation and plant growth promotion. The Virulence Factors Database (VFDB) was used to curate the virulence factor gene panel (29). AMR levels were quantified using the PATRIC3/mic prediction tool (67). A composite score was developed to characterize each bacterial isolate genome sequence with respect to the presence/absence of genes from the NF, PGP, and VF gene panels along with the predicted AMR levels. Details on the gene panels, AMR level, and the composite scoring system can be found in the Supplementary Methods.

### Experimental validation

Predictions made by computational phenotyping were validated using five distinct experimental assays: (1) Acetylene reduction assay for nitrogen fixation activity, (2) Phosphate solubilization assay, (3) Siderophore production assay, (4) Gibberellic acid production assay, and (5) Indole acetic acid production assay. Details of each experimental assay can be found in the Supplementary Methods.

## ACKNOWLEDGMENTS

We thank members of the INCAUCA Laboratory of Microorganismal Production for their support with the isolation of sugarcane-associated bacteria in Colombia. We thank the Kostka laboratory technicians Patrick Steck and Michael Blejwas for their support with laboratory analysis of sugarcane-associated bacteria at Georgia Tech.

